# A Chalcone Synthase-Like Bacterial Protein Catalyzes Heterocyclic C-Ring Cleavage of Naringenin to Alter Bioactivity Against Nuclear Receptors in Colonic Epithelial Cells

**DOI:** 10.1101/2022.04.22.489210

**Authors:** Ebru Ece Gülşan, Farrhin Nowshad, Meredith Davis Leigh, Jimmy W Crott, Stephen Safe, Arul Jayaraman, Kyongbum Lee

**Author notes:** E.E.G and F.N contributed equally to this work. Corresponding Authors: Kyongbum Lee, Department of Chemical and Biological Engineering, Tufts University, Medford, MA 02155. Arul Jayaraman, Artie McFerrin Department of Chemical Engineering, Texas A&M University, 3122 TAMU, College Station, TX 77843.

## Abstract

The human gut microbiota contributes enzymatic functions that are unavailable to host cells and play crucial roles in host metabolism, nutrient processing and regulating immune functions. As dietary compounds that are only partially absorbed, flavonoids are available for metabolism by gut microbiota, leading to diverse bioactive products. Combining prediction of enzyme promiscuity, metabolomics, and *in vitro* model systems, we identified a bacterial enzyme that can catalyze heterocyclic C-ring cleavage of naringenin. Culture experiments using a wild-type and mutant strain of *Bacillus subtilis* confirmed that the enzyme is a chalcone synthase-like polyketide synthase. The prediction-validation methodology developed in this work could be used to systematically characterize the products of gut bacterial flavonoid metabolism and identify the responsible enzymes and species. Further, we demonstrated that naringenin and its ring cleavage metabolites differentially engage the AhR and NR4A in intestinal epithelial cells. Our results suggest that the abundance of selected gut bacterial species impacts the profile of bioactive flavonoids and flavonoid-derived metabolites and thereby influences inflammatory responses in the intestine. These results are significant for understanding the mechanisms of gut microbiota-dependent effects of dietary flavonoids.

## Introduction

Flavonoids are the largest class of naturally occurring polyphenolic phytochemicals, numbering over 6,000 structurally distinct molecules (Panche et al., 2016).As dietary compounds, they are particularly abundant in plant-based foods that are considered health-promoting. The health benefits of flavonoids are generally associated with their antioxidant and anti-inflammatory activities (Ginwala et al., 2019), although the potency varies widely even among flavonoids of the same subclass. There is no clear consensus regarding the molecular mechanisms underlying the bioactivities of flavonoids. *In vitro* studies have shown that flavonoids can engage specific cellular receptors (Chatree et al., 2021; Zhang et al., 2003), but it remains to be established if and which of the pathways regulated by these receptors are responsible for the flavonoids’ beneficial effects *in vivo*. Further, it is unclear whether the flavonoids directly bind the receptors or are first transformed in the body into metabolic products which then activate the receptors.

In the intestine, potential molecular targets for flavonoids are the aryl hydrocarbon receptor (AhR) and nuclear receptor 4A (NR4A). Both are transcription factors capable of binding a variety of metabolite ligands to regulate inflammatory pathways in colonic epithelial cells (Lamas et al., 2018; Safe et al., 2016). Studies have shown that flavonoids can exhibit agonist or antagonist activities to induce or inhibit expression of AhR and NR4A target genes (Avior et al., 2013; Jin et al., 2018). These two receptors can also bind phenolic acids derived from flavonoids (Havlik et al., 2020; Kampa et al., 2004). In contrast to many flavonoid compounds, the phenolic acids are readily absorbed in the intestine and are typically found at higher plasma concentrations. For example, the average daily plasma concentrations of quercetin and naringenin in humans are on the order of 1 µM (Walle, 2004), whereas their major phenolic acid metabolites phenylacetic acid and phenylpropionic acid are present at hundred-fold or higher concentrations (Jenner et al., 2005). This raises the intriguing question whether flavonoid derived metabolites are quantitatively important ligands for the AhR, NR4A, and other flavonoid relevant receptors.

Upon ingestion, flavonoids are only partially absorbed and available for metabolism by commensal microorganisms residing in the intestine, or gut microbiota. Most dietary flavonoids (except for flavan-3-ols) are ingested in their glycoside form. The first step of their metabolism is deglycosylation. For *O*-coupled flavonoids, deglycosylation can be catalyzed by human glycosidases such as lactase-phlorizin hydrolase and cystolic β-glycosidase (Nemeth et al., 2003). *C*-glycosidases, however, are quantitatively metabolized by gut bacteria (Day et al., 2000). Gut bacterial enzymes are also required for hydrolysis of sugar moieties other than glucose. Following glycoside hydrolysis, the resulting aglycones can be absorbed and conjugated into glucuronide and sulfate forms in the intestine or liver by phase II enzymes. Alternatively, they can be further metabolized by the gut bacteria. Gut bacteria can perform a variety of reactions, including ring cleavage, de/methylation and di/hydroxylation, to generate metabolites that cannot be formed through host metabolism alone. In addition to providing an energy source, metabolism of flavonoids also serves as a detoxification mechanism for some gut bacterial species (25793210). Addition or removal of hydroxy and/or methoxy groups has been observed under anaerobic conditions at all three flavonoid rings (Braune and Blaut, 2011; Kim et al., 2014). Cleavage of a C-C bond, however, has only been reported for the central C-ring. Once a flavonoid has undergone C-ring cleavage, it can be further degraded into short-chain fatty acids (SCFAs) (from the A-ring) and hydroxyphenyl acids (from the B-ring).

Not all gut bacteria can catalyze the degradation of flavonoids into these metabolites. For example, only a handful of species have been identified that can perform *O*-demethylation of isoflavones and flavanones (Kim *et al*., 2014; Possemiers et al., 2005; Possemiers et al., 2008). Moreover, the enzymatic pathways of flavonoid metabolism in gut bacteria are largely unresolved. Flavonoids are not natural substrates of gut bacterial enzymes. Consequently, reactions of flavonoid metabolism have been attributed to more general classes of enzymes (Cragg et al., 2006). Modifications such as hydrolysis, de/methylation and di/hydroxylation have been linked to glycosyltransferases, methyltransferases and oxidoreductases. However, the specific enzymes that belong to these classes and the bacterial species carrying the enzymes remain to be elucidated.

Recent *in vitro* studies using human fecal cultures reported that *Eubacterium ramulus* and *Flavonifractor plautii* are capable of breaking down quercetin, luteolin and naringenin into SCFAs and phenolic acids (Braune et al., 2001; Schoefer et al., 2003). An earlier study by Miyake et al. showed that anaerobically grown *Clostridium butyricum* can cleave the C3-C4 bond in the C-ring of eriodictyol to form phenolic acids (Miyake et al., 1997). Pereira-Caro et al. co-incubated *Bifidobacterium longum* and *Lactobacillus rhamnosus* with orange juice flavanones under anaerobic conditions and identified several phenolic acids as metabolic products (Pereira-Caro et al., 2018). While these studies demonstrate that selected gut bacteria are capable of degrading flavonoids into C-ring cleavage products, the responsible enzymes are unknown.

In this study, we investigate the metabolism of a model flavonoid, naringenin, into its C-ring cleavage product, 3,4-hydroxyphenylpropanoic acid (34HPPA) by gut bacteria. Using metabolic modeling, we predict that this metabolism proceeds through a chalcone synthase-like bacterial polyketide synthase and confirm this prediction using *in vitro* culture experiments. Further, we demonstrate that naringenin and 34HPPA differentially engage the AhR and NR4A in intestinal epithelial cells, underscoring the impact of gut bacterial flavonoid metabolism on flavonoid bioactivity in the intestine. Prospectively, the prediction-validation methodology developed in this work could be used to systematically characterize the metabolism of flavonoid metabolites and identify the responsible enzymes and species, directly linking the formation of bioactive flavonoid metabolites to abundance of specific gut bacteria.

## Results

### Enzyme promiscuity-based model predicts that gut bacteria can metabolize diverse flavonoids

We adapted a tool for prediction of promiscuous enzyme activity on xenobiotic chemicals (PROXIMAL) (Yousofshahi et al., 2015) to investigate possible products of gut bacterial flavonoid metabolism. Previously, we assembled a model of metabolic reactions catalyzed by bacterial enzymes of murine intestinal microbiota based on operational taxonomic units (OTUs) detected in anaerobic batch culture of fecal material from 6-to 8-week-old female C57BL/6J mice (Lei et al., 2019). For the present study, we modified this model to more generally predict promiscuous enzyme activities in murine intestinal microbiota. To this end, the previous metabolic model was pared down to a subset of representative strains that overlap with the Mouse Intestinal Bacterial Collection (miBC) (Lagkouvardos et al., 2016). This yielded a total of 106 strains belonging to 18 genera. These strains were then searched against the KEGG GENOME database and UniProtKB to identify the enzymes encoded in their genomes and match these enzymes with corresponding Enzyme Commission (EC) numbers. The reactant-product (RCLASS) data associated with these EC numbers were analyzed using PROXIMAL to determine atom group transformations in the enzymes’ natural substrates. These atom group transformations defined operators for an enzyme that, when applied to potential nonnatural substrates having atom groups that match the enzyme’s natural substrate, predict the corresponding reaction products (Supplementary Figure S1). In total, the murine intestinal microbiota model enzymes mapped to approximately 12,000 unique atom group transformation operators.

These operators were applied to 19 dietary flavonoids (myricetin, gossypetin, quercetin, taxifolin, morin, robinetin, luteolin, kaempferol, fisetin, naringin, apigenin, baicalein, naringin, baicalein-*O*-7-glucoside, genistein, daidzein, daidzein-8-glucoside, genistein-8-glucoside, daidzein-7-glucoside) belonging to 5 major subclasses and their glucoside forms found in fruits and vegetables (Supplementary Table S1). Metabolism of exogenous chemicals often occurs in two steps, with the first and second steps resulting in substrate activation and metabolism, respectively. To mimic this process, the products predicted by the operators for each of the 19 flavonoids were used as substrates for a second round of predictions. Combined, the two rounds predicted 1,427 metabolites having unique KEGG compound identifiers. We did not attempt an exhaustive search of the published literature for these compounds, as the lack of reported findings regarding gut bacterial metabolism of a flavonoid does not necessarily indicate that the metabolism does not occur. When gut bacterial metabolism of a flavonoid was reported, however, the experimental evidence largely agreed with our predictions. For example, 12 of the 16 predicted naringenin metabolites were detected in cultures of individual gut bacteria or fecal material incubated with naringenin (Braune and Blaut, 2016; Chen et al., 2019a; Xiao and Lee, 2017; Zou et al., 2015).

We next evaluated the results from our enzyme promiscuity based predictions against two other computational tools for predicting small molecule metabolism, BioTransformer (Djoumbou-Feunang et al., 2019) and Way2Drug (Rudik et al., 2016). This analysis focused on five flavonoids representing the major subclasses flavanol (quercetin), flavanone (naringenin), flavone (apigenin and luteolin), and isoflavone (genistein). The analysis also included naringin, the glucoside form of naringenin. The predictions by computational tools were compared for products of methylation, hydroxylation, dihydroxylation, hydrogenation, dehydrogenation, C-ring cleavage, and hydrolysis. Overall, our enzyme promiscuity based approach (PROXIMAL) generated the most comprehensive coverage of reaction products, except for hydrogenation products (Figure 1). Way2Drug predicted the largest number of products for hydrogenation, but the fewest for dihydroxylation and C-ring cleavage. PROXIMAL and Way2Drug predicted the same methylation and dehydrogenation products, which were larger than those predicted by BioTransformer. All three tools predicted the same set of hydrolysis products.

**Figure 1.**
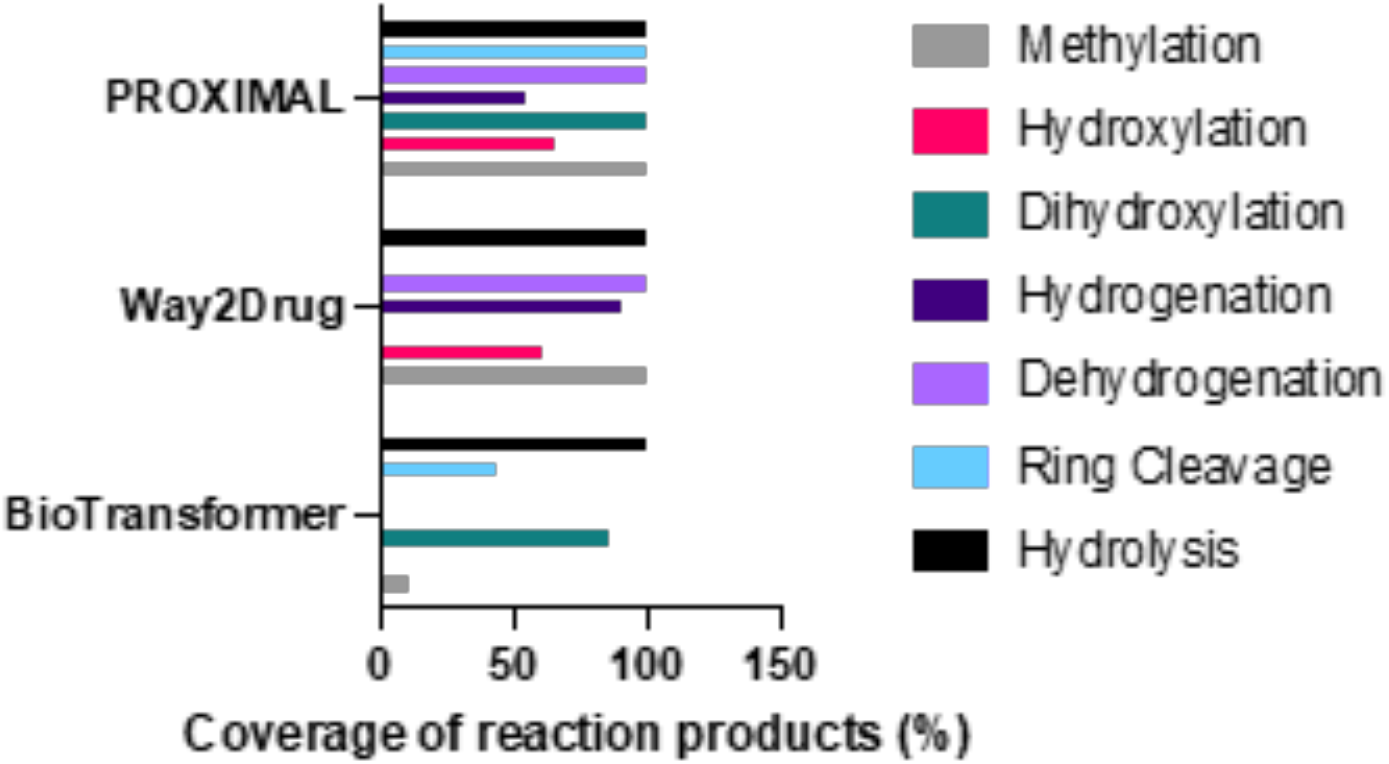
Comparison of reaction products predicted by PROXIMAL, Way2Drug, and BioTransformer for five flavonoids representing the major subclasses of flavanol (quercetin), flavanone (naringenin), flavone (apigenin and luteolin), isoflavone (genistein). Coverage was calculated with respect to the total number of distinct reaction products collectively predicted by the three tools. Full (100%) coverage by a tool for a reaction type indicates that the tool predicted all metabolites predicted by the other two tools for the same reaction type.

### Gut bacterial metabolism of flavonoids varies by molecular structure and taxa

We investigated the impact of flavonoid structure on the predicted reactions. Structural similarity scores were calculated for 15 aglycones using the SIMCOMP2 tool (Hattori et al., 2010). Figure 2A shows a multidimensional scaling (MDS) map where the distance between a pair of compounds corresponds to the dissimilarity (1 – similarity score) between the pair. Overall, flavonoids from the same subclass were structurally more similar to each other than compounds from other subclasses and grouped closer together on the MDS plot. For example, almost all the flavonols (fisetin, myricetin, morin, robinetin, quercetin, and gossypetin) grouped together in the top half of the plot except for kaempferol. The isoflavones (genistein and daidzein) formed a tight pair in the bottom of the plot. However, the flavones did not show this grouping; luteolin and baicalein grouped with the flavonols, whereas apigenin grouped closer to the isoflavones. These trends showed that the subclass of a flavonoid only partially predicts the structural similarity to other flavonoids. We observed a similar trend when we constructed an MDS map based on the flavonoids’ similarity of predicted reactions (Figure 2B). The isoflavones and flavonols formed two groups on the left and right side of the plot, respectively, whereas the flavones did not show this grouping. A correlation analysis confirmed that there was a modest, but significant association between the structural and reaction dissimilarities of flavonoids (Figure 2C). Based on these results, we chose quercetin, luteolin, naringenin, and genistein for further analysis to represent flavonoids of different subclasses having varying structures and predicted reaction profiles. We also included apigenin, which belongs to the same subclass as luteolin but grouped more closely with the isoflavones and naringenin, a flavanone.

**Figure 2.**
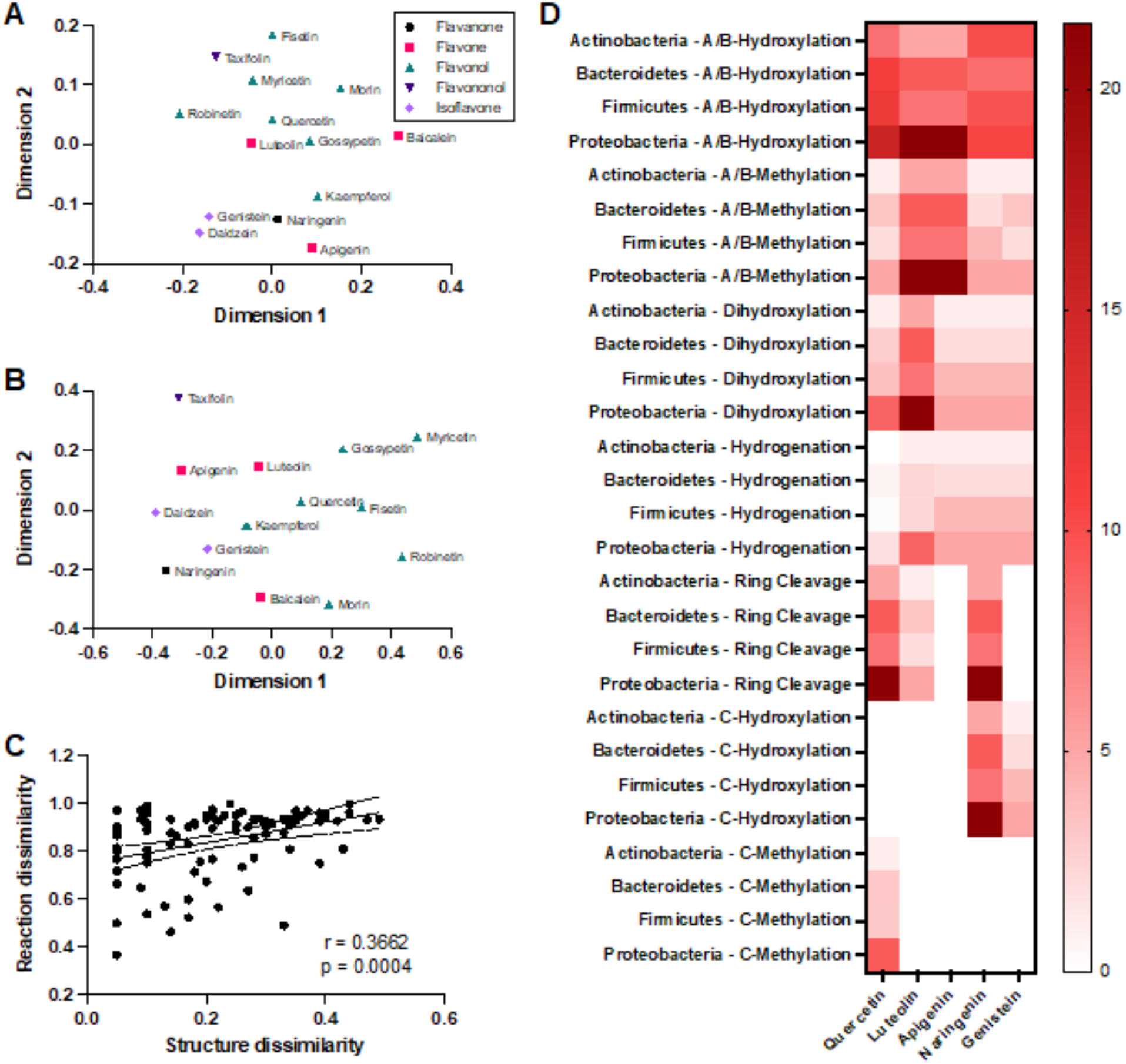
Impact of flavonoid subclass and structure on metabolism predictions across reaction types and bacterial taxa. (A) Multidimensional scaling (MDS) map for 15 flavonoid aglycones, where the compounds’ coordinates were assigned based on a matrix of relative pairwise distances representing structural dissimilarities calculated using the SIMCOMP2 tool. Symbols and colors indicate the compounds’ subclasses. (B) MDS map where the compounds’ coordinates were assigned based on their predicted reaction patterns. (C) Correlation between structural and reaction dissimilarities of flavonoids. Solid and dashed lines show the best fit linear regression model and 95% confidence intervals, respectively. (D) Distribution of predicted flavonoid metabolizing enzymes across different bacterial phyla. Color scale corresponds to the number of matching enzymes in the phylum. The number of matches for a phylum was normalized by the number of strains included in the model for the phylum. Each row corresponds to a different combination of phylum and type of reaction.

We next investigated the distribution of predicted flavonoid metabolizing enzymes among the model’s bacterial groups. We tabulated the enzymes corresponding to the PROXIMAL operators for predicted metabolites of the selected flavonoids and searched for matching enzymes in the microbiota model. The matches were determined based on the first three digits of the enzymes’ EC numbers. We did not use the fourth digit, which specifies the natural substrate or cofactor of an enzyme. A central premise of our predictions is that enzymes can catalyze reactions of non-natural substrates. We focused on methylation, hydroxylation, dihydroxlation, hydrogenation and C-ring cleavage as these reactions were most frequently predicted for the selected flavonoids. To investigate if the functional group position matters, we separately tabulated hydroxylation and methylation reactions on A/B and C rings. Figure 2D shows the predicted distribution of flavonoid metabolizing enzymes across different bacterial groups at the phylum level. The trends shown in the heatmap were similar at lower taxonomic levels, as genera and families belonging to the same phylum had similar sets of enzymes predicted to metabolize flavonoids (Supplementary Figure S1). Hydroxylation and methylation on A and B rings were predicted for all phyla and flavonoids. Naringenin and genistein had the largest number of enzymes predicted to catalyze A or B ring hydroxylation, while apigenin and luteolin had the most matches for A or B ring methylation. Dihydroxylation followed a similar trend, with luteolin having the largest number of matching enzymes. Hydrogenation was predicted to occur less commonly for all flavonoids, with Actinobacteria lacking the enzymes altogether. Unlike A and B ring reactions, C ring reactions were predicted for only some of the flavonoids. Hydroxylation was predicted mainly for naringenin, whereas methylation was predicted only for quercetin. C-ring cleavage showed a heterogenous distribution across flavonoids and phyla. While we did not predict any enzymes that could catalyze this reaction for apigenin and genistein, we predicted multiple such enzymes for quercetin and naringenin in strains from all four phyla.

### Dose- and time-dependent increase in 3-(4-hydroxyphenyl)-propionic acid concentrations in the monocultures

Enzymatic cleavage of a C-C bond in the C-ring of flavanones has been extensively studied, yet *F. plautii* is the only species detected in murine intestine reported to catalyze this reaction. The results in Figure 2D suggested that there are other murine gut bacteria that can catalyze this reaction. To experimentally validate this prediction, we cultured selected species from the microbiota model in the presence of naringenin and analyzed the spent medium and cell pellets for accumulation of 34HPPA, a metabolite that results from C-ring cleavage of the flavanone. We observed a dose- and time-dependent increase in 34HPPA when anaerobically grown *F. plautii* were treated with naringenin (Figure 3A), consistent with previously reported experiments (Schoefer *et al*., 2003; Stevens et al., 2019). We observed a time-dependent increase in 34HPPA when naringenin was added to a culture of *E. coli*, but this accumulation occurred independent of naringenin dose (Figure 3B). This indicated that *E. coli* can produce 34HPPA but does not use naringenin as a major substrate. The amount of 34HPPA detected in the *E. coli* culture was significantly higher than *F. plautii*. This is likely due to the higher growth rate of *E. coli*, as the amounts are comparable for the two cultures when normalized to the culture OD (data not shown). Neither *L. plantarum* nor *P. lactis* culture showed a dose- and time-dependent accumulation of 34HPPA upon naringenin treatment (Figures 3C&D). These results are consistent with our prediction that species in these two genera cannot catalyze naringenin C-ring cleavage (Supplemental Table S2).

**Figure 3.**
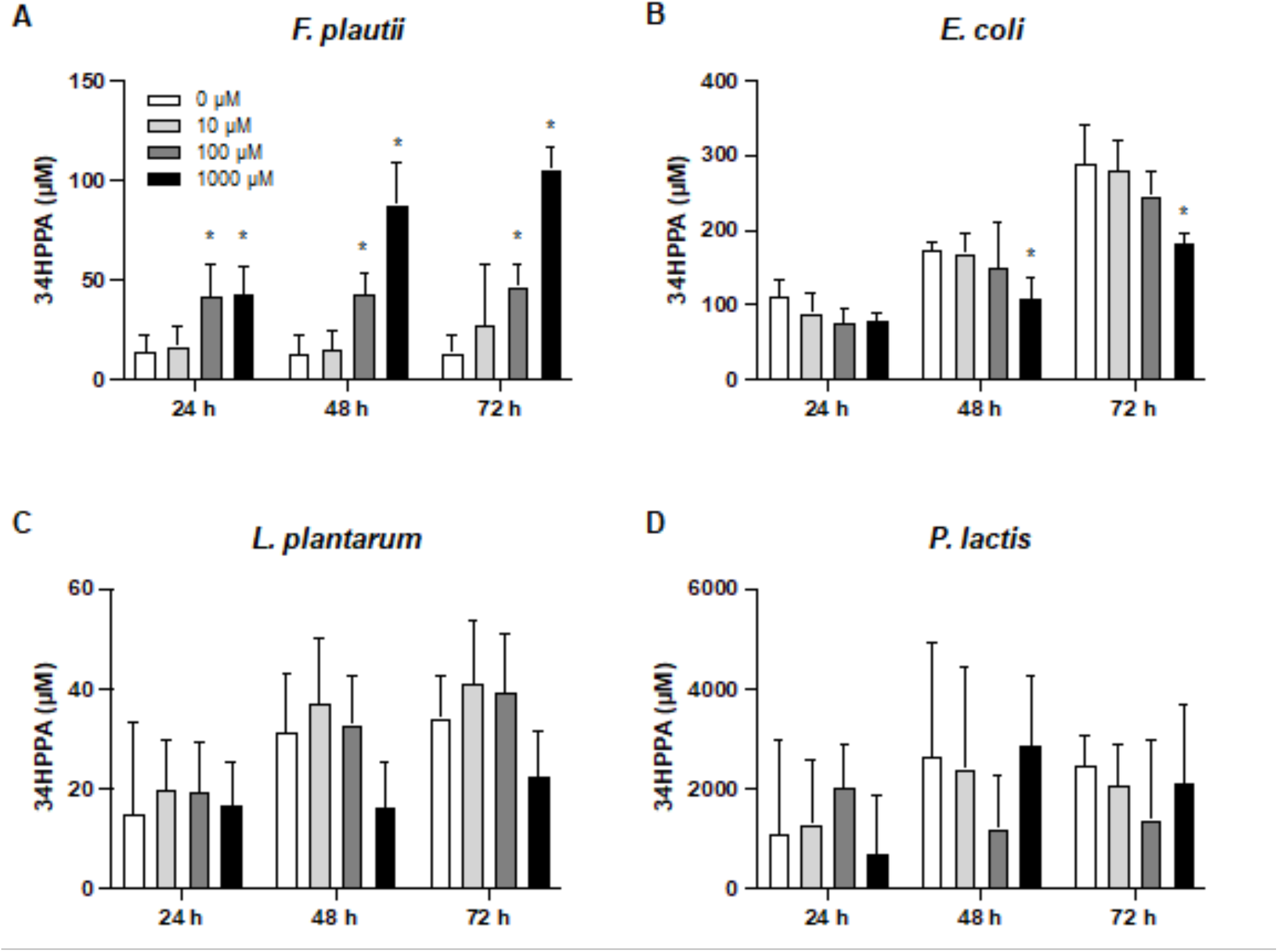
Concentrations of 3-(4-hydroxyphenyl) propionic acid (34HPPA) in naringenin treated cultures of (A) *F. plautii*, (B) *E. coli*, (C) *L. plantarum*, and (D) *P. lactis*. Data shown are means ± SD (N = 3 biological replicates). Asterisk (*) indicates significant difference (p < 0.05) compared to the vehicle control (0 µM) at the corresponding time point.

We next investigated the enzyme(s) responsible for naringenin C-ring cleavage. In plants, flavanone C-ring cleavage is catalyzed by chalcone isomerase (CHI). However, none of the strains in the microbiota model has a gene for CHI or a CHI ortholog. We further examined the PROXIMAL results by clustering the predicted flavonoid reactions based on their RCLASS similarity. This analysis found that a chalcone synthase (CHS)-like enzyme could accept naringenin as a substrate to form naringenin chalcone. In plants, CHS has been shown to bind naringenin at a highly conserved active site (Ferrer et al., 1999b). To explore the possibility that a CHS-like enzyme catalyzes naringenin chalcone formation in gut bacteria, we created a phylogenetic tree with all bacterial strains in the microbiota model that have genes homologous to CHS from *Medicago sativa* (a legume expressing CHS with strong naringenin binding activity) with a sequence similarity score > 80%. This analysis found that *B. subtilis* has a bacterial polyketide synthase (*bcsA*) with a high degree of sequence similarity to *M. sativa* CHS.

To investigate if this enzyme catalyzed naringenin C-ring cleavage in *B. subtilis*, we incubated wild-type *B. subtilis* with varying concentrations of naringenin (10, 100 and 1000 µM), and observed a dose- and time-dependent increase in 34HPPA (Figure 4, orange bars). We performed the same dose-response experiment using a mutant strain, *B. subtilis* BKK22050, that lacks *bcsA*. Unlike the wild-type strain, the mutant strain did not show a dose- and time-dependent increase in 34HPPA (Figure 4, blue bars), indicating that CHS activity is necessary for naringenin C-ring cleavage in *B. subtilis*.

**Figure 4.**
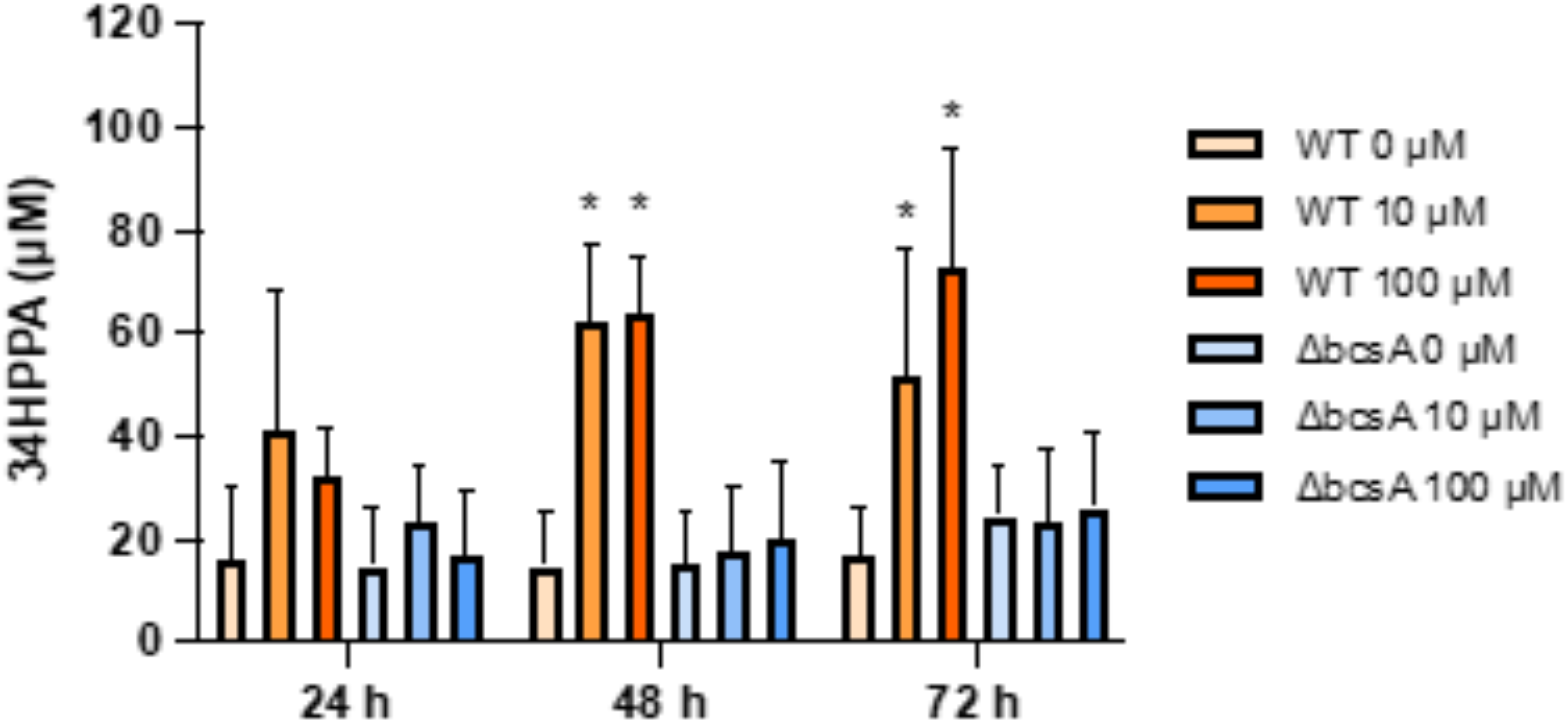
Concentrations of 3-(4-hydroxyphenyl) propionic acid (34HPPA) in naringenin treated cultures of wild-type (WT, orange bars) and mutant (blue bars) *B. subtilis*. Data shown are means ± SD (N = 3 biological replicates). Asterisk (*) indicates significant difference (p < 0.05) compared to the vehicle control (0 µM naringenin) at the corresponding time point.

### Model fecal culture microorganisms metabolize naringenin through C-ring cleavage

To investigate naringenin degradation (Figure 5A) in a mixed community of gut bacteria, we incubated murine fecal cultures with varying doses of naringenin. At the highest dose (100 mM), we observed a significant ∼40% decrease in naringenin after 48 h of incubation, indicating net utilization of naringenin (Figure 5B). Although high concentrations of 34HPPA were detected in the culture, this was not dependent on naringenin dose (data not shown). One possible explanation for this is that the fecal culture microorganisms produced 34HPPA from other sources as we observed for *E. coli* (Figure 3B). We also detected phloretin (Figure 5C), which lies upstream of 34HPPA in the C-ring cleavage pathway of naringenin, but the fecal culture concentrations were below the quantification limit of our LC-MS assay (∼10 µM).

**Figure 5.**
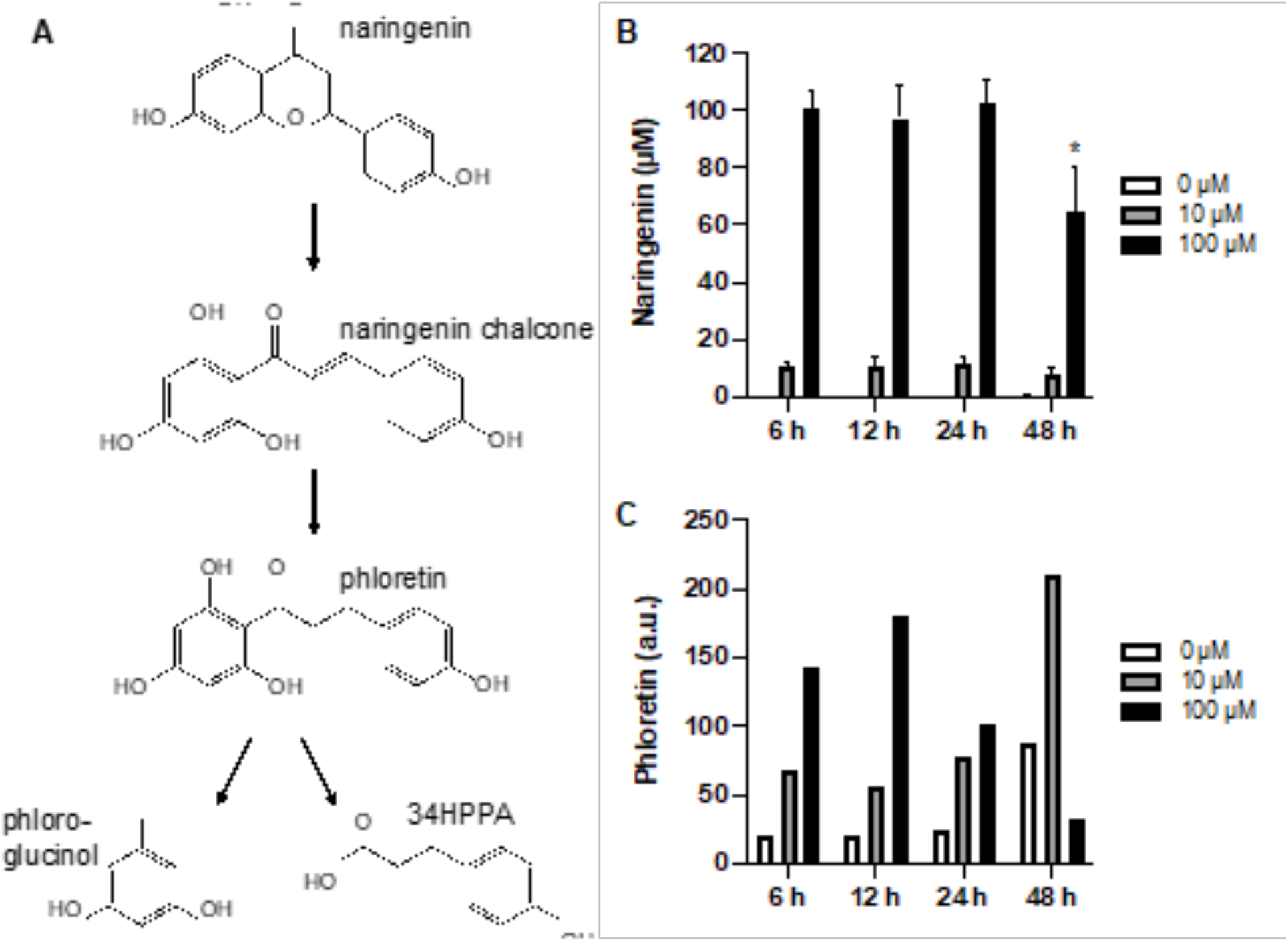
Naringenin and phloretin concentrations in fecal cultures treated with naringenin. (A) Proposed pathway for naringenin metabolism via C-ring cleavage in the fecal culture. Concentrations of (B) naringenin and (C) phloretin in the fecal culture as a function of naringenin dose. Data shown are means ± SD (N = 3 biological replicates). Asterisk (*) indicates significant difference compared to initial time point. (p < 0.05).

To determine if conversion of phloretin to 34HPPA occurred in the fecal culture, we supplemented the culture medium with ^13^C_6_-phloretin (but not naringenin) and monitored the accumulation of ^13^C_6_-34HPPA. We observed a time-dependent decrease in ^13^C_6_-phloretin (Figure 6A) and concomitant increase in ^13^C_6_-34HPPA (Figure 6B). We also detected high concentrations of unlabeled 34HPPA (Figure 6C), consistent with the monoculture trends indicating that murine gut bacteria produce 34HPPA not only through flavonoid metabolism, but also other pathways.

**Figure 6.** Conversion of phloretin-(hydroxyphenyl-^13^C_6_) to ^13^C_6_-34HPPA in fecal culture. Concentrations of (A) phloretin-(hydroxyphenyl-^13^C_6_), (B) ^13^C_6_-34HPPA, (C) and unlabeled 34HPPA. Data shown are means ± SD (N = 3 biological replicates).

### Naringenin and its metabolites elicit different biological activities

Having confirmed that naringenin can be metabolized by gut bacteria to 34HPPA by way of naringenin chalcone and phloretin, we next investigated if the parent flavanone and its metabolic products elicit different biological responses. We have previously shown that many flavonoids have AhR ligand activity (Jin *et al*., 2018; Park et al., 2019; Safe et al., 2021). Therefore, we assayed for induction of CYP1A1, CYP1B1, and UGT1A1 in murine (YAMC) as well as human (Caco-2) colonic epithelial cell models. In Caco-2 cells, naringenin had no effect and its chalcone had minimal effect on CYP1A1 and CYP1B1 gene expression (Figure 7A). Unlike naringenin, its chalcone dose-dependently induced UGT1A1 gene expression. Results in YAMCs followed a similar trend, with naringenin chalcone eliciting significant dose-dependent induction of CYP1B1 and UGT1A1 expression and naringenin showing weaker activity (Figures 7E&F).

**Figure 7.**
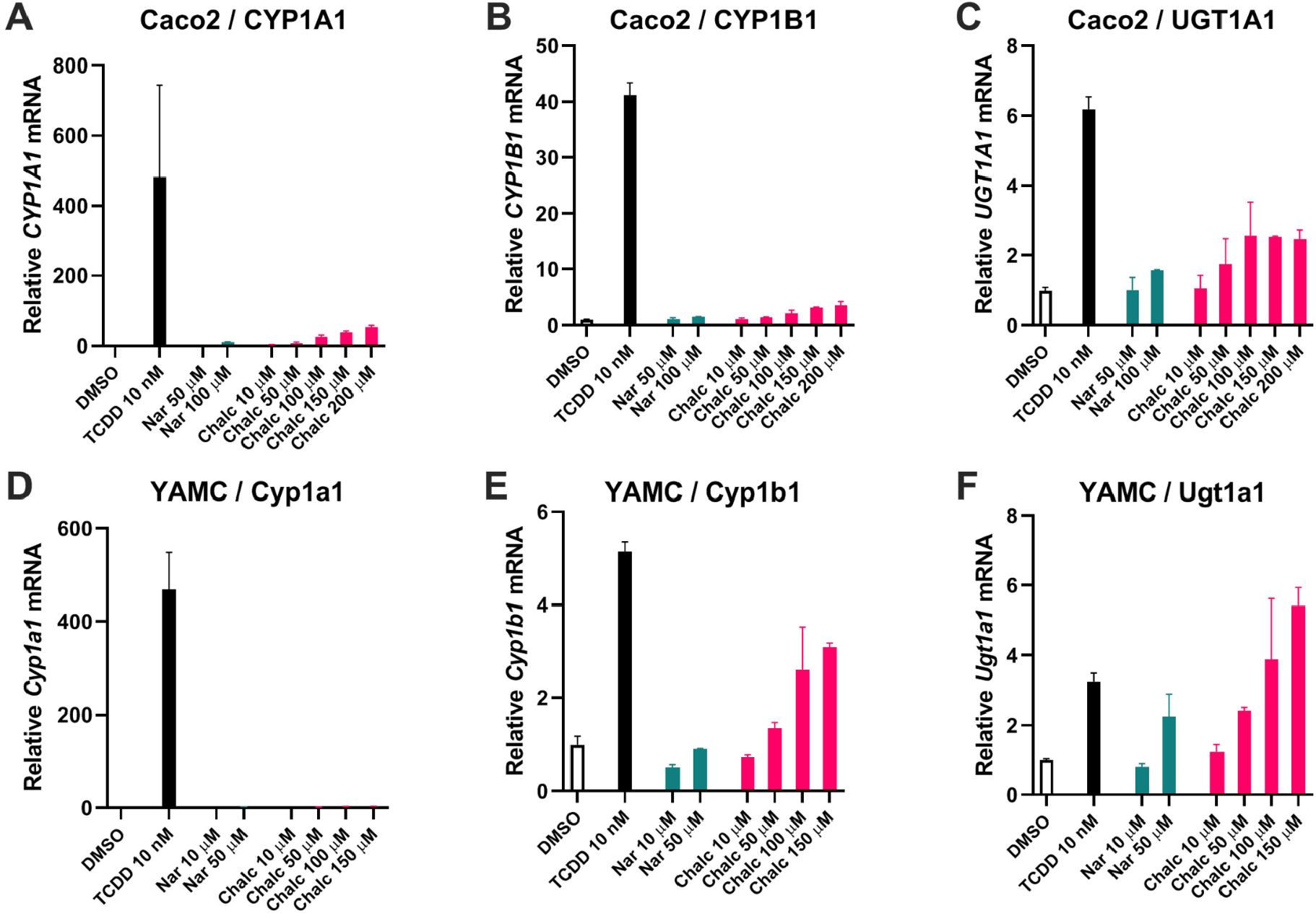
Induction of AhR target enzymes CYP1A1, CYP1B1, and UGT1A1 by naringenin (Nar), naringenin chalcone (Nar Chalc), and phloretin in (A-C) Caco-2 cells and (D-F) YAMCs. The cells were treated with varying concentrations of the indicated metabolites for 24 h. Data shown are means ± SD (N = 3 biological replicates). Asterisk (*) indicates significant difference (p < 0.05) compared to vehicle control (DMSO).

One of the factors influencing AhR ligand activation in colonic epithelial cells is crosstalk with other small molecule gene expression modulators. Recent studies showed that short-chain fatty acids (SCFAs) such as acetate, butyrate and propionate can enhance TCDD-induced Ah-responsive gene expression in colonic epithelial cells (Park *et al*., 2019). We therefore investigated if 34HPPA, which, unlike the chalcone intermediate, did not induce AhR target gene expression (data not shown), could influence gene expression in conjunction with an agonist. In Caco2 cells, 34HPPA significantly enhanced UGT1A1 and CYP1B1 expression when combined with TCDD. This effect was similar to butyrate, which is known to have a synergistic effect on AhR ligand induced CYP1A1, CYP1B1 and UGT1A1 gene expression in these cells (Figure 8).

**Figure 8.**
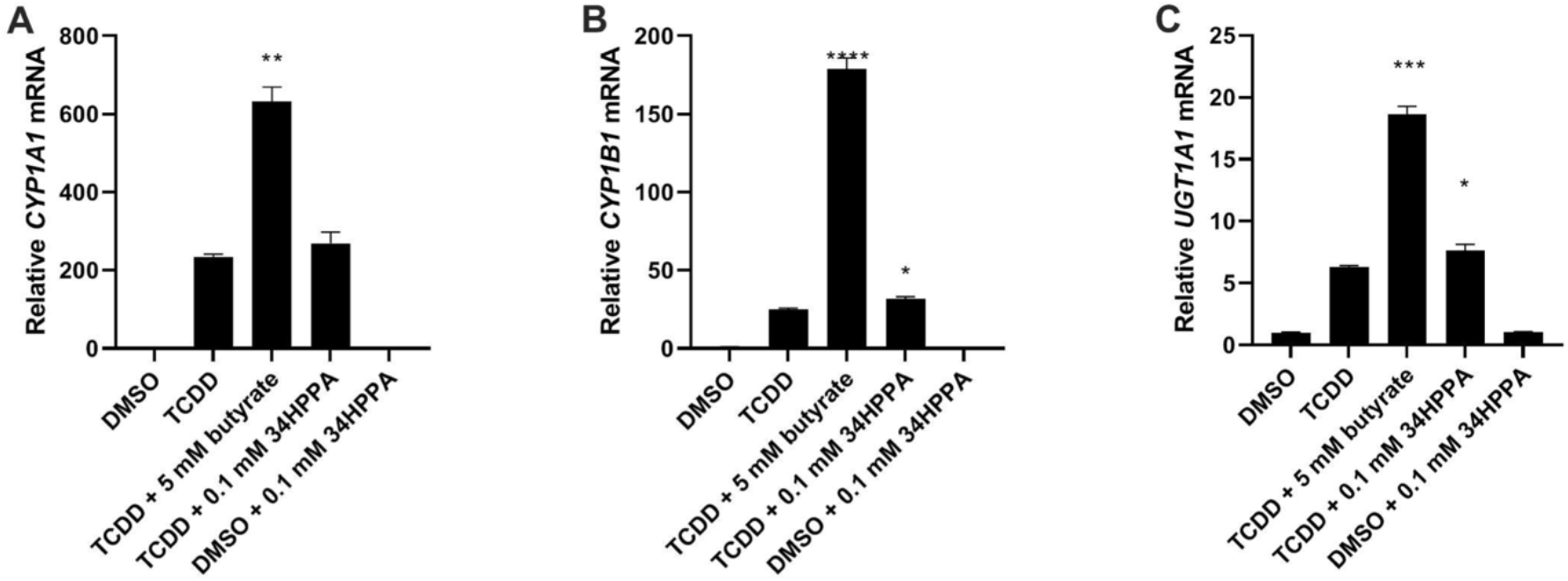
Induction of AhR target enzymes (A) CYP1A1, (B) CYP1B1 and (C) UGT1A1 by 34HPPA alone and in combination with 10 nM TCDD. Data shown are means ± SD (N = 3 biological replicates). Asterisk (*) indicates significant difference (p < 0.05) compared to TCDD.

There is increasing evidence that the nuclear receptor 4A subfamily, in addition to the AhR, could also play a role in mediating the biological activities of flavonoids. Recent *in vitro* studies have demonstrated that kaempferol and quercetin can bind the ligand binding domain (LBD) of NR4A1 to inhibit pro-oncogenic NR4A1-regulated pathways (Shrestha et al., 2021). We investigated if naringenin or its gut bacterial metabolites can also bind nuclear receptor 4A LBDs. Direct binding assays using LBDs of NR4A1 and NR4A2 showed that naringenin has strong affinity for both LBDs (Figures 9A&B) with K_D_ values of 4.5 and 5.2 mM, respectively. In contrast, 34HPPA did not show any binding activity for either LBD (Figures 9C&D).

**Figure 9.**
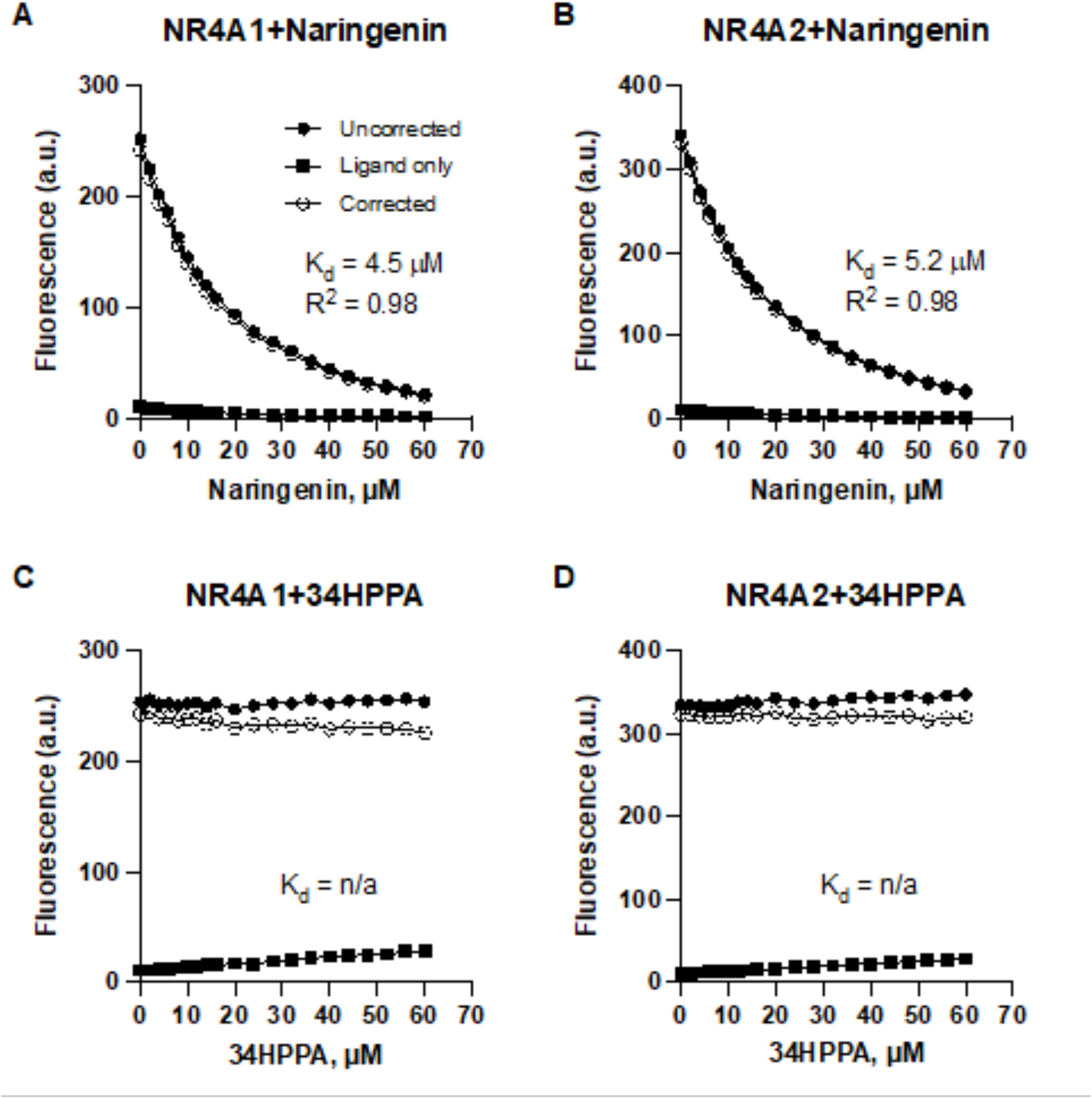
Concentration-dependent quenching of tryptophan fluorescence in the ligand-binding domain (LBD) of NR4A1 and NR4A2 with (A, B) naringenin and (C, D) 34HPPA.

Finally, having found that naringenin and its microbial metabolic products elicit different biological responses as AhR active compounds and NR4A1/2 ligands, we next investigated their immunomodulatory effects in colonic epithelial cells by stimulating Caco-2 cells with interleukin-1b (IL-1b) and measuring the secretion of IL-8, a neutrophil chemotactic factor. Whereas naringenin dose-dependently decreased IL-8 secretion, neither 34HPPA nor phloretin had a significant effect (Figure 10)

**Figure 10.**
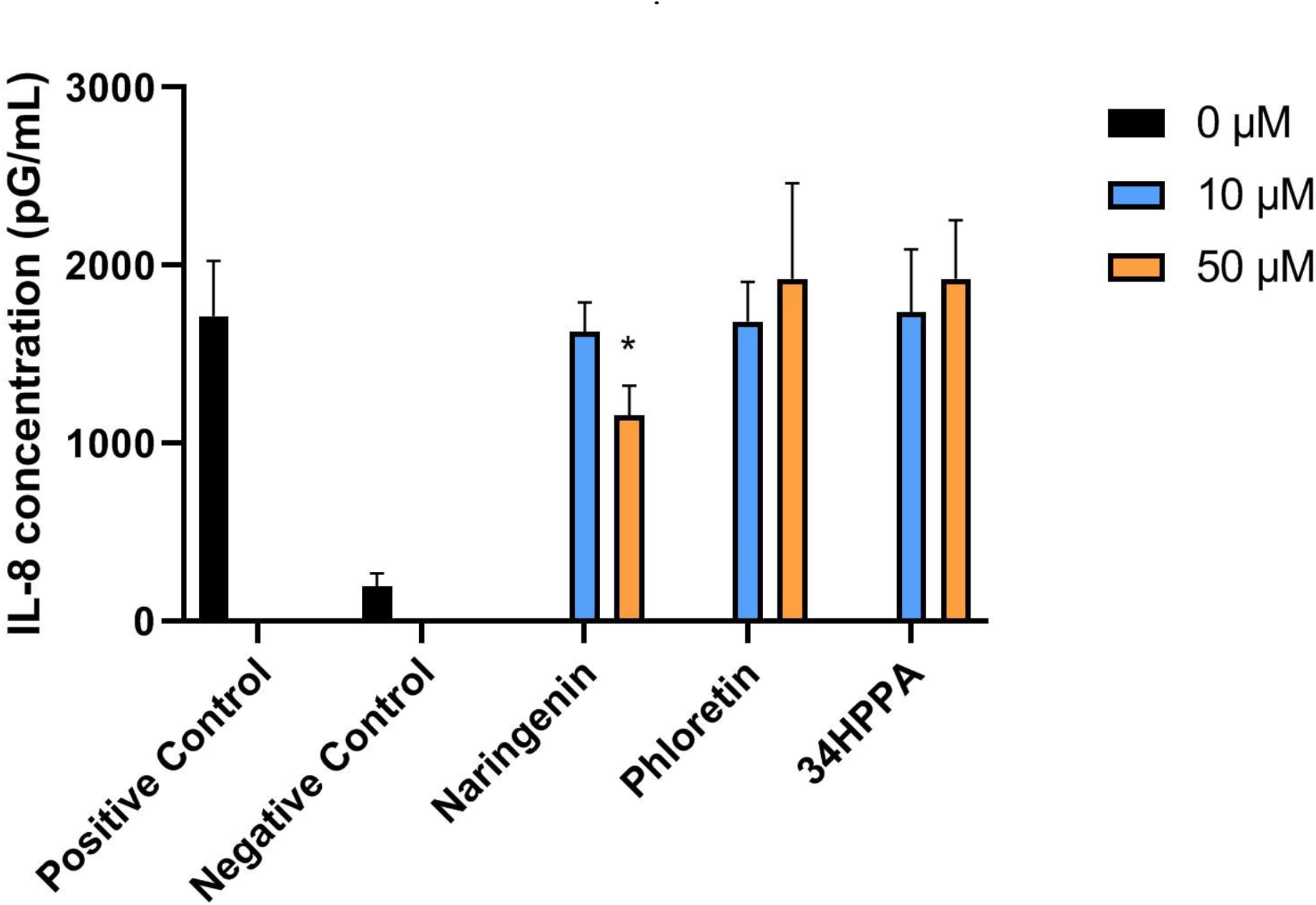
IL-8 concentration in conditioned medium following IL-1β stimulation of Caco-2 cells treated with naringenin, 34HPPA or phloretin. Positive and negative controls indicate cells that were stimulated with and without IL-1β in the absence of flavonoids, respectively. Data shown are means ± SD (N = 3 biological replicates). Asterisk (*) indicates significant difference (p < 0.05) compared to positive control.

## Discussion

In this work, we utilized a prediction tool for promiscuous enzyme activity on nonnatural substrates to investigate gut bacterial metabolism of dietary flavonoids. We identified a novel bacterial enzyme that can catalyze the C-ring cleavage reaction of a flavanone and showed that a degradation pathway that proceeds through this reaction is active in anaerobic batch culture of murine fecal microbiota. Finally, we showed that intermediates of naringenin’s C-ring cleavage degradation pathway elicit different biological activities as AhR and NR4A1/2 ligands and mediate differential immunomodulatory responses in intestinal epithelial cell lines.

Evaluation of the enzyme promiscuity-based predictions against *in vitro* and *in vivo* studies reported in the literature found strong corroborating evidence for the predicted metabolites. Among 16 predicted naringenin metabolites, 12 were detected in cultures of individual gut bacteria or fecal material incubated with naringenin (Braune and Blaut, 2016; Chen *et al*., 2019a; Xiao and Lee, 2017; Zou *et al*., 2015). Burapan et al. reported methylation products of luteolin and apigenin by *Blautia sp*. MRG-PMF1 under anaerobic conditions that match the predictions of the present study (Burapan et al., 2017). Isorhamnetin and tamarixetin were detected in rat plasma as methylated metabolites of quercetin (Claudine et al., 1996; Mohos et al., 2019), consistent with our prediction results. Using fecal slurry cultures from human donors, Di Pede et al. found that quercetin degrades into various phenylpropanoic, benzoic and phenylacetic acid derivatives (Di Pede et al., 2020). These metabolites were expected to result from C-ring cleavage of quercetin, which was predicted for all four major bacterial phyla by our enzyme promiscuity model (Figure 3D).

Not all predictions could be corroborated with reported findings. For example, Soukup et al. (Soukup et al., 2021) and Lee et al. (Lee et al., 2017) suggested that the first step of anaerobic microbial degradation of genistein is hydrogenation on C-ring, a reaction our model did not predict. Further, we did not find evidence in the literature that genistein undergoes any of the other predicted reactions such as de/methylation and di/hydroxylation. It is possible that certain flavonoid derived metabolites may not be readily measured *in vitro* due to challenges in culturing the microorganisms that express the responsible enzymes. Additionally, the resulting metabolites may be intermediates that are further degraded into smaller organic acids. Another reason some metabolites were not observed by previous studies could be that these metabolites were not in the scope of the studies’ targeted analyses. In this regard, computational predictions of promiscuous enzyme activity complement experimental investigations in systematically characterizing the metabolism of flavonoids, which are not well-defined natural substrates of gut bacteria.

There have been limited attempts to develop prediction methods for gut microbial metabolism of dietary phytochemicals, whereas there are several well-known approaches for predicting the metabolism of chemicals in the environment or human tissue, such as enviPath (Wicker et al., 2016) and Meteor Nexus (Greene et al., 1999). These approaches have typically used rule-based methods to generate biotransformation pathways, where the rules are derived from a knowledgebase of enzymatic reactions. BioTransformer is a recently developed tool that combines knowledge-based and machine learning approaches to generate biotransformation rules (Djoumbou-Feunang *et al*., 2019). Notably, this tool includes a module for gut microbial metabolism built from databases of metabolites detected in bodily fluids after consumption of polyphenols (Neveu et al., 2010) and other dietary phytochemicals (https://phytohub.eu). One key difference between our method and BioTransformer is that the latter’s knowledgebase, which includes a fixed set of gut microbial enzymes, models a generic gut microbiome rather than a particular microbiome having user-defined composition. Another difference is that BioTransformer assigns many of its rules for gut microbial metabolism to unspecified enzymes, whereas the present study links each biotransformation operator to one or more specific gut microbial enzymes. One advantage of rule-based methods such as BioTransformer is that it can predict multi-step transformations. While the PROXIMAL algorithm could in principle be used for this purpose by processing the outputs from one iteration as inputs for the next iterations, the result would be a combinatorial problem, as the algorithm is not paired with a reasoning engine to select reactions that are more likely to occur. In this regard, a future implementation of PROXIMAL could incorporate learning-based approaches from retrosynthesis, where a heuristic search algorithm (e.g., Monte Carlo Tree Search) is used to explore the biotransformation space (Koch et al., 2020).

In addition to BioTransformer, we evaluated our predictions against another tool, Way2Drug RA, which identifies the reacting atoms of drug-like compounds undergoing biotransformation by human cytochrome P450 (CYP) enzymes and transferases. Notably, its reaction database includes microbial flavonoid reactions such as aromatic hydroxylation and *O*-glucuronidation. This reaction database, however, cannot customized by the user to model specific microbiomes. Compared to both BioTransformer and Way2Drug RA, the PROXIMAL implementation performed in the present study on a murine gut microbiota model yielded a more comprehensive set of predictions, while also modeling a specific set of promiscuous enzymatic reactions that could occur in a user-defined microbiome. The latter feature is critical for investigating which gut bacteria are responsible for flavonoid metabolism. In the present study, we successfully validated a novel prediction that *B. subtilis* can metabolize a flavanone via C-ring cleavage.

Naringenin is a naturally occurring phytochemical that belongs to the flavonoid subclass flavanone and is abundant especially in citrus fruits. The starter unit for naringenin biosynthesis is 4-coumaroyl-CoA, which is derived from either phenylalanine or tyrosine depending on the plant (Álvarez-Álvarez et al., 2015; Kyndt et al., 2002). Chalcone synthase (CHS) combines 4-coumaroyl-CoA with three malonyl-CoA molecules to produce naringenin chalcone, which is then rapidly isomerized to naringenin by chalcone isomerase (CHI) (Sun et al., 2015). This reaction is reversible, and CHI can also cleave the C-ring of naringenin to produce naringenin chalcone (Supplementary Figure S2).

**Supplementary Figure S2.**
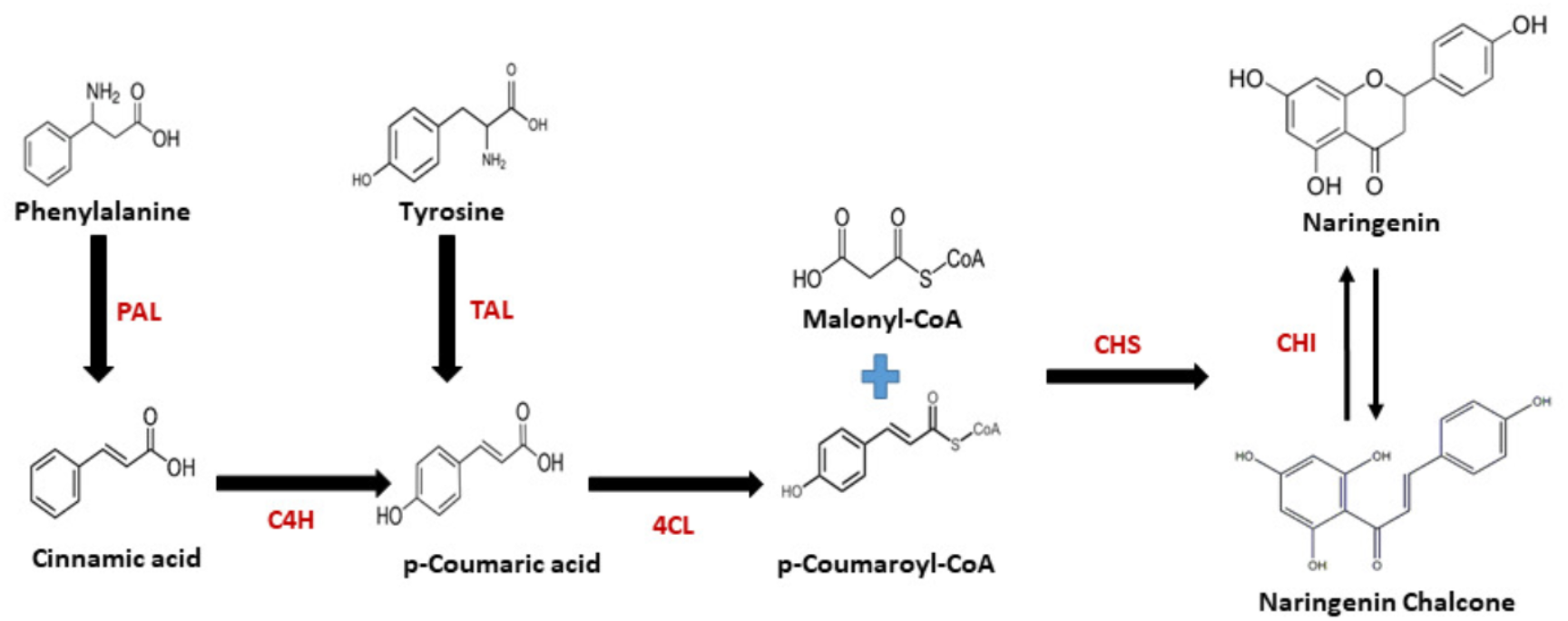
Naringenin biosynthesis pathway in plants. PAL: Phenylalanine ammonia-lyase, TAL: Tyrosine ammonia-lyase, C4H: Cinnamate-4-hydroxylase, 4CL: P-coumarate-CoA ligase, CHS: Chalcone synthase, CHI: Chalcone isomerase.

Gut bacterial degradation of flavonoids results in the formation of various organic acids, notably hydroxylated phenylpropionic acids. Although it is well-established that this degradation proceeds through cleavage of the heterocyclic C-ring (Booth et al., 1956), the species and key enzymes involved remain largely unknown. In the present study, we have identified a bacterial CHS-like enzyme that can catalyze C-ring cleavage of the flavanone naringenin. Enzymes of the CHS superfamily share high similarity in their amino acid sequence, structure and general catalytic principles and contain the conserved Cys-His-Asn catalytic triad at the binding site (Sun *et al*., 2015). Although 4-coumaroyl-CoA is the natural substrate for plant CHS, earlier studies have confirmed that CHS of *Medicago sativa* can also use naringenin as a substrate (Ferrer et al., 1999a). *In vitro* monocultures of *F. plautii* and *B*.*subtilis* show a dose- and time-dependent accumulation of 34HPPA, whereas *E. coli* and other gut bacteria predicted to lack the CHS-like enzyme did not show this trend. However, we did not detect other naringenin ring cleavage pathway intermediates such as naringenin chalcone and phloretin in the F. plautii and B. subtilis monocultures. This could be due to rapid metabolism of these intermediates into 34HPPA. We also did not detect phloroglucinol, another product of the ring cleavage pathway, which can be completely degraded by intestinal microbiota into acetate, butyrate and CO_2_ (Possemiers et al., 2011).

Unlike the monocultures of *F. plautii* and wild-type *B. subtilis*, we did not detect a dose-dependent increase in 34HPPA in the fecal cultures upon naringenin addition. While 34HPPA is a major product of naringenin metabolism by gut bacteria (Chen et al., 2019b), the fecal culture environment likely contained other sources for this phenolic acid. Aromatic amino acids such as-phenylalanine or tyrosine, which are present in the GMM used for the fecal culture experiments, are also potential precursors for 34HPPA. This could explain the presence of the higher concentration of 34HPPA in fecal cultures independent of naringenin dose and time. A more specific intermediate of naringenin metabolism via C-ring cleavage is phloretin, which was detected at elevated levels in naringenin treated fecal cultures relative to vehicle treated controls. However, the phloretin concentrations were below quantifiable levels. To more sensitively trace phloretin’s metabolic fate, we repeated the experiment by supplementing the culture medium with ^13^C_6_-phloretin instead of naringenin. We observed a time-dependent decrease in ^13^C_6_-phloretin concentration and concomitant concentration increase in ^13^C_6_-34HPPA. Our results also show a dose-dependent increase in phloretin 6 and 12 h after naringenin treatment, but not at 24 and 48 h. One possible explanation is that accumulation of phloretin leads to the activation of downstream enzymes that can utilize phloretin. Further, we did not detect any unlabeled phloretin in the culture, indicating that the phloretin detected in the naringenin treatment experiment is derived from naringenin rather than any other medium component. Taken together, these results suggest that phloretin derived from naringenin is metabolized to 34HPPA in the fecal culture. On the other hand, the conversion of phloretin to 34HPPA is low. Whereas labeled phloretin concentration decreased to ∼30 mM by 72 h, labeled 34HPPA increased to only ∼20 mM. It is possible that 34HPPA was further degraded to other phenolic acids, e.g., 4-hydroxyphenyl acetic acid, 3-phenyl propionic acid, and benzoic acid. Another possibility is that phloretin was converted to phloridzin (phloretin 2′-*O*-glucose), which can be formed by bacterial phloretin-2′-*O*-glycosyltransferase (P2′GT) in presence of glucose in bacteria culture media (Zhang et al., 2016).

Recent structure activity relationship experiments on flavonoids in host cells indicated that there is a significant difference in AhR ligand activity between the parent compounds and their gut bacterial metabolites (Blay et al., 2018; Koper et al., 2019). For example, quercetin is a weak agonist/partial antagonist for AhR in T47D breast cancer cells (19286049). In the same cell line, 3,4-dihydroxyphenylacetic acid (DOPAC), a microbial metabolite derived from quercetin (Havlik *et al*., 2020), did not affect AhR activation (Kampa *et al*., 2004). Flavonoid metabolism depends on microbiota composition. Rechner et al. did not detect DOPAC as a product of quercetin metabolism in an *in vitro* mixed culture model of human colonic microbiota. Instead, they reported 3-hydroxyphenylacetic acid and 3-(3-hydroxyphenyl)-propionic acid as the major quercetin metabolites (Rechner et al., 2004). Tamura et al. studied quercetin metabolism by fecal microbiota from healthy elderly human subjects and found that Fusobacteriaceae and Enterobacteriaceae affect quercetin bioavailability by inhibiting its degradation by other bacteria. They also reported that the abundance of Sutterellaceae and Enterobacteriaceae are positively correlated with quercetin degradation, suggesting that the fate of quercetin depends on microbiota composition (Tamura et al., 2017). Thus, responsible enzymes are likely unevenly distributed across taxa, which leads to variations in metabolism and the immunomodulatory effects.

In this study, we characterized naringenin and its ring cleavage metabolites as AhR ligands in Caco-2 and YAMC cells. The results from gene expression assays indicate that naringenin chalcone has moderate AhR activity, whereas naringenin and the downstream metabolites have little activity. We also found YAMCs were more responsive than Caco-2 cells to induction of AhR gene expression by naringenin chalcone. Our results agree with other studies on ligand activation of the AhR that the cell type plays a significant role in target gene expression (Zhang *et al*., 2003).

Another factor influencing AhR ligand activation is crosstalk with other gene expression modulators. Studies have shown that short-chain fatty acids (SCFAs) enhance the responsiveness of the AhR to structurally diverse ligands due, in part, to their role as histone deacetylase inhibitors (Jin et al., 2017). Park et al., reported that acetate, butyrate, and propionate enhance TCDD-induced CYP1A1 expression in Caco-2 cells (Park *et al*., 2019). In the present study, we found that 34HPPA exhibited SCFA-like synergy with an AhR agonist, significantly enhancing TCDD-induced CYP1A1 and CYP1B1 expressions even though the naringenin metabolite by itself did not show AhR ligand activity. These results suggest that gut bacterial metabolism of dietary flavonoids could influence AhR-mediated responses in the intestine not only by shaping the profile of AhR active flavonoid ligands, but also by producing other small molecules that interact with AhR pathways. While we demonstrated that 34HPPA is the major metabolite of naringenin degradation via C-ring cleavage, the fecal culture environment contains aromatic amino acids such as phenylalanine or tyrosine that are also precursors for 34HPPA. Therefore, the 34HPPA concentration in the intestine is likely high even when flavonoids are only partially degraded. Our results also suggest naringenin chalcone is an AhR-active ligand. Given that naringenin, naringenin chalcone and 34HPPA are likely to occur together in the intestine, it is important to consider the potential for synergistic activation of the AhR by 34HPPA, naringenin chalcone, and other microbially derived AhR ligands. To the best of our knowledge, this is the first study to investigate the synergistic induction of AhR-regulated gene expression by a microbially derived flavonoid metabolite.

NR4A is a family of orphan nuclear receptors that plays a significant role in innate and adaptive immunity, maintaining cellular homeostasis and in pathophysiology. In many diseases, cellular stressors induce various NR4A members. For example, NR4A1 levels increase in many solid tumor-derived cancer cells with enhanced metabolic rates (Safe and Karki, 2021). In this study, we showed that naringenin binds with high affinity to NR4A1 and NR4A2, whereas its metabolite 34HPPA was inactive. In effect, gut microbial metabolism of naringenin leads to the inactivation of an NR4A1/2-active compound.

Having found that naringenin and its metabolites elicit different biological responses as AhR and NR4A ligands, we investigated whether these differences extend to their immunomodulatory effects. We observed significant dose-dependent decrease in IL-1ß induced IL-8 production in Caco-2 cells upon naringenin treatment, whereas phloretin and 34HPPA had no effect. Hamers et al. reported that overexpression of NR4A1 contributes to the anti-inflammatory response of Caco-2 cells by decreasing the expression of IL-8 (Hamers et al., 2015). This is consistent with the findings from the present study, which show that the NR4A1/2-active compound naringenin inhibits IL-8 expression in Caco-2 cells, while 34HPPA is NR4A1/2-inactive and does not elicit the anti-inflammatory effect.

In summary, this study presents a prediction-validation methodology that could be used to systematically characterize gut microbial flavonoid metabolism and identify the responsible microorganisms and enzymes. We report on novel experimental evidence that a CHS-like bacterial polyketide synthase can catalyze heterocyclic C-ring cleavage of naringenin. Our model predicts that this family of enzymes is available in only a subset of gut bacteria, which could explain variations in dietary flavonoid metabolism observed in both mice and humans with different intestinal microbiomes. We further showed that a flavonoid and its metabolites elicit different biological activities as AhR and NR4A1/2 ligands, which could explain their differential effects in mediating inflammatory responses in intestinal epithelial cells.

## Materials and Methods

### Cell lines, Bacterial Strains, Culture Conditions, and Reagents

The young adult mouse colonic (YAMC) cell line was originally obtained from the R. H. Whitehead Ludwig Cancer Institute (Melbourne, Australia). The cells were maintained in RPMI 1640 medium with 5% fetal bovine serum, 5 units/ml mouse interferon-γ (IF005) (EMD Millipore, MA), 0.1% ITS “−” minus (insulin, transferrin, selenium) (41-400-045) (Life Technologies, Grand Island, NY) at 33 °C in the presence of 5% CO2. Caco-2 human colon cancer cells were obtained from the American Type Culture Collection (ATCC, Manassas, VA) and maintained in Dulbecco’s modified Eagle’s medium (DMEM) nutrient mixture supplemented with 20% (v/v) fetal bovine serum (FBS), 10 mL/L 100× MEM nonessential amino acid solution (Gibco), and 10 mL/L 100× penicillin-streptomycin (Pen-Strep) solution (Sigma-Aldrich, St. Louis MO) at 37°C in the presence of 5% CO2 and the solvent (dimethyl sulfoxide, DMSO) used in the experiments was ≤ 0.2%. TCDD (99%) used in this study was synthesized in the Safe laboratory. *B. subtilis* ATCC 23857, *F. plautii* ATCC 49531, *L*.*plantarum* ATCC 14917, and *E*.*coli* ATCC BAA-1430 were obtained from ATCC (Manassas,VA). *P. lactis* DSM 28626 was obtained from the German Collection of Microorganisms and Cell Cultures (DSMZ, Braunschweig, Germany). *B. subtilis* BKK22050 was obtained from the Bacillus Genetic Stock Center (BGSC, Columbus, OH). Both wild type (ATCC 23857) and mutant (BKK22050) strains of *B. subtilis* were cultured in nutrient broth supplemented with potato extract (Sigma-Aldrich, St. Louis MO). *L. plantarum* was cultured in MRS broth (Sigma-Aldrich, St. Louis MO). *P. lactis* was cultured in Luria-Bertani (LB) broth (Sigma-Aldrich, St. Louis MO). *E. coli* and *F. plautii* were cultured in brain heart infusion (BHI) broth (Becton, Dickinson and Company, Franklin Lakes, NJ) supplemented with 0.5% yeast extract (Becton, Dickinson and Company, Franklin Lakes, NJ), 0.05% cysteine (Alfa Aesar, Tewksbury, MA), 0.05% hemin, and 0.02% vitamin K (Sigma-Aldrich, St. Louis MO).

### Bacterial monoculture

The *B. subtilis* strains were cultured aerobically in nutrient broth as described above. Cultures grown to an OD (absorbance at 600 nm) of 1.0 were diluted 10-fold and treated with varying doses of naringenin (10, 100, or 1000 μM, stock solutions were prepared using DMSO as solvent and then filter sterilized) or vehicle (0.1% v/v DMSO). Following the naringenin or vehicle treatment, *B. subtilis* cultures were incubated at 37 °C for 72 h. *P. lactis, L. plantarum, E. coli*, and *F. plautii* cultures were grown in their respective media as mentioned above inside an anaerobic chamber with 85% N_2_, 10% CO_2_, 5% H_2_ (Coy Lab, Grass Lake, MI) and treated with naringenin or vehicle in a similar manner. Each monoculture experiment was run in triplicate. Samples were collected for metabolite extraction by sacrificing the cultures after 24, 48 and 72 h of incubation.

### Fecal culture

Fresh fecal material from 14-week-old female C57BL/6J mice were harvested and immediately transported to an anaerobic chamber with 85% N_2_, 10% CO_2_, 5% H_2_ (Coy Lab, Grass Lake, MI) using anaerobic transport medium (BD Diagnostics, Franklin Lakes, NJ). All reagents were introduced into the anaerobic chamber a day before the experiment to ensure that they were pre-reduced by the start of the experiment. 10 mL of anoxic phosphate-buffered saline (PBS) containing 0.5g/L of cysteine was added to 1 g of fecal sample. The suspension was homogenized by vigorous shaking and vortexing. Samples were left for 15 min to allow large particles to settle. To culture fecal bacteria, gut microbiota medium or GMM was used which was prepared as described previously (Goodman et al., 2011). Sterile conical tubes containing 9 mL of GMM were inoculated with 300 µL of the fecal slurry. The inoculated tubes were then treated with varying doses of naringenin (10 or 100 μM) or vehicle (0.1% v/v DMSO) at the time of inoculation (day 0). The fecal cultures were then incubated under anaerobic condition at 37 °C and sacrificed for metabolite extraction after 6, 12, 24 and 48 h of incubation. For the ^13^C isotopic labeling experiment, the fecal cultures were treated with 100 μM of phloretin-(hydroxyphenyl-^13^C_6_) at the time of inoculation and incubated anaerobically as described above. The cultures were sacrificed for metabolite extraction after 24, 48, and 72 h of incubation.

### Metabolite Extraction

Metabolites were extracted from the harvested cell culture samples using a solvent-based method. Briefly, 50 µL of culture sample was mixed with 200 µL of ice-cold methanol using a vortex mixer and kept on ice for 60 minutes. The sample-solvent mixture was then centrifuged at 15,000g at 4 °C for 5 min. The supernatant was filtered through a 0.22 µm sterile nylon filter (Corning). The filtrate was then passed through a 3 kDa molecular weight cutoff filter (Amicon Ultra, Sigma-Aldrich) placed in a collection tube by centrifuging the sample tube at 15,000g for 45 min at 4 °C. The filtrate from the collection tube was then mixed with 2 mL sterile water, lyophilized to a pellet using a freeze dryer (FreeZone, Labconco Corporation) and stored at -80 °C until further analysis. Prior to LC-MS/MS analysis, the freeze dried samples were reconstituted in 250 µL of 50% (v/v) methanol/water.

### Targeted Analysis of Naringenin Metabolites

The extracted samples were analyzed for naringenin, phloretin, and 34HPPA using product ion scan experiments performed on a triple-quadrupole time-of-flight instrument (5600+, AB Sciex) coupled to a binary pump high-performance liquid chromatography (HPLC) system (1260 Infinity, Agilent). Chromatographic separation was achieved on a Synergi 4 µm Hydro, 250 mm × 2 mm, 80 A reverse phase column (Phenomenex, Torrance, CA) maintained at 30 °C using a solvent gradient method. Solvent A was 0.2% (v/v) formic acid in water and solvent B was 100% acetonitrile. The gradient method was as follows: 0–30 min (95% A to 45% A), 30–38 min (45% A to 95% A). The flow rate was set to 0.6 mL/min. The injection volume was 10 µL. Raw data were processed in PeakView 2.1 to determine the ion peaks. The peak areas in the extracted ion chromatograms were manually integrated using MultiQuant 2.1 (AB Sciex). The identities of detected compounds were confirmed by matching their retention time and/or MS/MS spectra to high-purity standards run on the same instrument using the same method.

### AhR Activity Assay

Caco-2 and YAMC cells were maintained in their respective media as described above. Cells were seeded at 60–70% density and were sub-confluent after treatment and subsequent analysis. YAMC cells (1.2 × 10^7^ cells) were treated with TCDD (10 nM) and/or compounds such as naringenin (50 - 100 µM) and naringenin chalcone (10 – 200 µM) for 2 or 24 h. Caco-2 cells (5 × 10^6^ cells) were treated with TCDD and/or compounds for 2 h. For co-treatment, Caco-2 cells were treated with 100 µM of 34HPPA as well as 5 mM butyrate (Thermo Fisher Scientific, Inc. Waltham, MA) for 24 hours. The cells were then fixed with 1% formaldehyde, and the cross-linking reaction was stopped by addition of 0.125 M glycine. After washing with phosphate-buffered saline, cells were scraped and pelleted.

### Quantitative Real-Time Reverse Transcriptase PCR

Total RNA was extracted using an RNA isolation kit from the cells according to the manufacturer’s protocol. cDNA synthesis was performed from the total RNA of cells using the High-Capacity RNA-to-cDNA Kit (Applied Biosystems, Foster City, CA). Real-Time PCR was carried out using Bio-Rad SYSR Universal premix for 1 min at 95 °C for initial denaturing, followed by 40 cycles of 95 °C for 15 s and 60 °C for 1 min in the Bio-Rad iCycler (MyiQ2) real-time PCR System. Gene expression values were analyzed using the comparative CT method and normalized to expression levels of TATA-binding protein (TBP). The sequences of the primers used for real-time PCR were as follows: human ***CYP1A1*** sense 5′-GAC CAC AAC CAC CAA GAA C-3′, antisense 5′-AGC GAA GAA TAG GGA TGA AG-3′; ***CYP1B1*** sense 5′-GGA TAT CAG CCA CGA CGA AT-3′, antisense 5′- ATT ATC TGG GCA AAG CAA CG-3′; ***UGT1A1*** sense 5′-GAA TCA ACT GCC TTC ACC AAA AT-3′, antisense 5′-AGA GAA AAC CAC AAT TCC ATG TTC T-3′; ***TBP*** sense 5′ -GAT CAG AAC AAC AGC CTG CC- 3′, antisense 5′ -TTC TGA ATA GGC TGT GGG GT- 3′. Mouse ***Cyp1a1*** sense 5′-CAG GAG AGC TGG CCC TTT A-3′, antisense 5′-TAA GCC TGC TC ATC CTG TG-3′; ***Cyp1b1*** sense 5′- GGA TAT CAG CCA CGA CGA AT -3′, antisense 5′- ATT ATC TGG GCA AAG CAA CG -3′; ***Ugt1a1*** sense 5′- ATG GCT TTC TTC TCC GGA AT- 3′, antisense 5′- TCA GAA AAA GCC CCT ATC CC -3′; ***TBP*** sense 5′ - GAA CAA TCC AGA CTA GCA GCA - 3′, antisense 5′ - GGG AAC TTC ACA TCA CAG CTC - 3′.

### NR4A1/2 Binding Assay

Binding affinities of naringenin and 34HPPA for NR4A1 and NR4A2 were assayed based on quenching of tryptophan fluorescence as previously described (Shrestha *et al*., 2021). The ligand binding domains (LBD) of NR4A1 and NR4A2 were incubated at a fixed concentration (0.5 μM) in PBS with different concentrations of naringenin and 34HPPA. Fluorescence spectra were obtained at 285/330 nm (slit width 5 nm).

### Cytokine Measurement

Caco-2 cells were cultured in DMEM supplemented with 10% (v/v) FBS, 2 mM glutamine, and 1% penicillin–streptomycin (Life Technologies, Grand Island, NY). Sub-confluent cultures were maintained in a humidified incubator at 37°C with 5% CO_2_. Cells were seeded in a 96-well plate at a density of 1.5 × 10^4^ cells per well and allowed to attach overnight. The next day, the growth medium was replaced with serum-free medium. The cells were then treated with naringenin, 34HPPA, or vehicle for 2 h followed by addition of IL-1β (50 ng/mL). After 24 h, the supernatants were collected and clarified using centrifugation at 1500 RPM and 4 °C for 10 min. Secretion of IL-8 was measured in the cell-free supernatant samples using the Human IL-8/CXCL8 Quantikine ELISA Kit, following the manufacturer’s protocol (R&D Systems, Minneapolis, MN, D8000C).

### Murine Intestinal Microbiota Model

To define the atom group transformation operators used for predicting flavonoid metabolism, we assembled a model of metabolic reactions catalyzed by enzymes in representative strains of murine intestinal microbiota. We previously compiled a list of organisms to represent bacteria detected in anaerobic batch culture of cecal contents from 6-to 8-week-old female C57BL/6J mice (Lei *et al*., 2019). We compared this list against the Mouse Intestinal Bacterial Collection (miBC) (Lagkouvardos *et al*., 2016) and removed strains belonging to genera absent in miBC. The rationale for this step was to obtain a reduced list of organisms that are not specific for the experimental system used in the previous study and more broadly representative of culturable murine intestinal bacteria. We then added miBC strains absent from the reduced list but belong to a genus represented in the list. These steps yielded a total of 106 strains belonging to 18 genera common to miBC and the cecal culture. These strains were processed as described previously (Lei *et al*., 2019) to obtain a matrix of organisms and their enzymes’ EC numbers.

### Prediction of Flavonoid Metabolism

To predict products of flavonoid metabolism and gut bacterial enzymes responsible for these products, we modified a scheme previously developed to analyze the products of xenobiotic transformation reactions catalyzed by liver cytochrome P450 enzymes (Yousofshahi *et al*., 2015). In this scheme, the reactant-product pair(s) (RPAIR) of an enzymatic reaction (Moriya et al., 2010) are analyzed to identify changes in the reactant’s atom group(s) that transform the reactant into the product. These changes define the enzyme’s transformation pattern that is associated with a reaction center and its first- and second-level neighboring atoms. If a substrate of interest has an atom group that matches this pattern for an enzyme, then a transformation operator associated with the enzyme is applied to generate a product from the substrate. In the present study, we performed the RPAIR analysis on enzymes in the above-described murine intestinal microbiota model. A total of 5,932 unique atom group transformation operators were identified for the model. These operators were applied to each of the 19 flavonoid compounds to predict their bacterial metabolites. To allow for two-step transformations analogous to phase I and II reactions for xenobiotic transformation, the products from one round of metabolite predictions were used as inputs for a second round. The metabolites from both rounds were filtered to eliminate trivial products such as simple sugars, CO_2_, and water and compounds that are not cataloged as metabolites in the KEGG Compound database.

### Predictions of Flavonoid Metabolizing Enzymes and Microorganism

Enzymes in the microbiota model corresponding to operators that generate C-ring cleavage metabolites for naringenin were assembled into a list with their corresponding RPAIRs. These RPAIRs were clustered using a binning method based on atom-pair descriptors generated from their SMILES identifiers in ChemMine Tools (Backman et al., 2011). The similarity of RPAIRs was calculated as the Tanimoto coefficient of corresponding atom-pair descriptors. Distances between clusters were determined by single linkage. The compounds which fall into the same group with naringenin were examined further. If the reaction resembles the validated transformation, the reported enzymes were marked due to their potential to catalyze the transformation. Each reaction was assumed to be reversible unless specified otherwise. Predicted key enzymes were matched with the organism by using the organism-enzyme matrix from the murine cecal microbiota model. The matches were validated from available genome annotation databases. (Shrestha *et al*., 2021)

