## Supplementary figures and images for "A Chalcone Synthase-Like Bacterial Protein Catalyzes Heterocyclic C-Ring Cleavage of Naringenin to Alter Bioactivity Against Nuclear Receptors in Colonic Epithelial Cells"

### Supplemental Figure S1

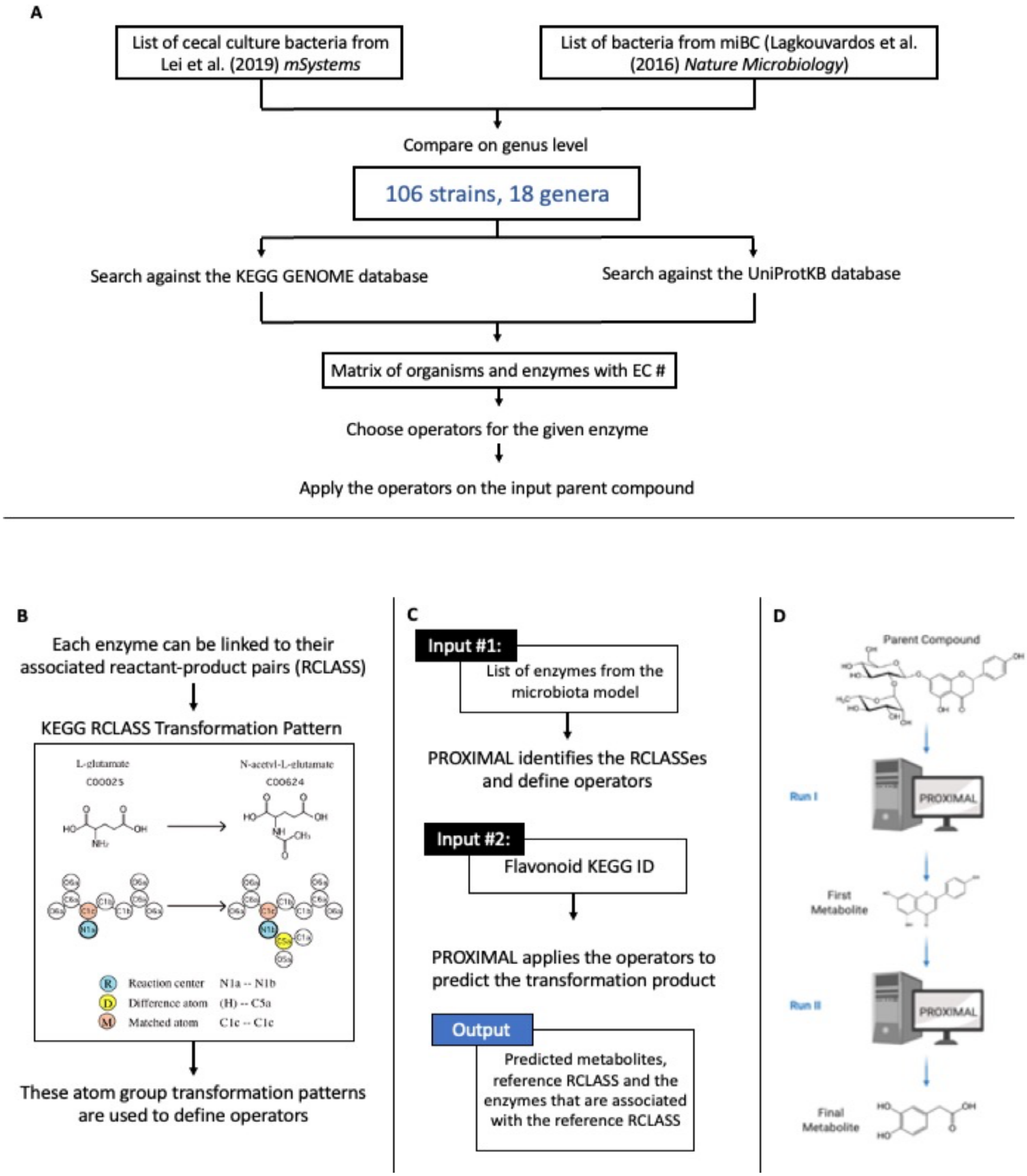
